# An Immunocompetent Mouse Model of Degarelix Induced Chemical Castration to Investigate Acquired Resistance to Androgen Deprivation

**DOI:** 10.64898/2026.10.07.757377

**Authors:** Amanda Robotham, Diane Costanzo-Garvey, Grace Waldron, Leah M Cook, Grinu Mathew

**Affiliations:** Eppley Institute for Research in Cancer & Allied Diseases, University of Nebraska Medical Center, Omaha, NE, USA; Nebraska Center for Molecular Target Discovery and Development, University of Nebraska Medical Center, Omaha, NE, USA; Fred & Pamela Buffett Cancer Center, University of Nebraska Medical Center, Omaha, NE, USA; Center for Cancer Research, National Cancer Institute, Bethesda, MD, USA

**Keywords:** Prostate cancer, chemical castration, Degarelix, androgen deprivation therapy, immunocompetent mouse model, acquired resistance to ADT

## Abstract

Prostate cancer (PC) remains a significant health concern as the second leading cause of cancer-related death in American men. Androgen deprivation therapies represent the standard of care for patients with PC, regardless of disease stage. Gonadotropin-releasing hormone (GnRH) antagonists, such as Degarelix, are commonly used hormone therapies for PC and work by suppressing testosterone production. Previous studies indicate a prostate involution-regeneration cycle following surgical castration and testosterone addback. However, whether a similar response occurs following pharmacologic androgen deprivation with agents such as Degarelix remains poorly understood. In particular, the temporal dynamics of prostate involution and subsequent regeneration after chemical castration have not been systematically defined. Our study utilizes immunocompetent mouse models to evaluate the systemic and temporal effects of Degarelix treatment and determine its efficacy in suppressing PC tumor growth. We show that continuous Degarelix treatment significantly suppressed systemic testosterone levels in mice, whereas testosterone levels returned to physiological baseline following Degarelix withdrawal. Gross anatomical, H&E, and Trichrome analyses demonstrated that Degarelix induced castration resulted in prostate involution, characterized by reduced gland size and increased stromal remodeling. This phenotype was reversed after Degarelix withdrawal, highlighting the reversibility of pharmacological androgen deprivation. Immunofluorescence staining revealed changes in basal and luminal epithelial cell populations within the prostatic gland, indicating changes in prostate glandular architecture following Degarelix induced castration and subsequent regeneration. Next, we evaluated the impact of chemical castration on PC tumor growth. Although Degarelix initially suppressed tumor growth, this response was not sustained through the study endpoint, suggesting the emergence of acquired resistance to androgen deprivation. These findings establish this immunocompetent model as a physiologically relevant platform for dissecting the mechanisms that drive therapeutic resistance and tumor progression following androgen deprivation. Our findings highlight the effectiveness of chemical castration with Degarelix and establish a robust model to study both therapeutic responses and the emergence of castration resistance in PC.

## Introduction

Prostate cancer (PC) continues to pose a significant health concern, being the most diagnosed cancer and the second leading cause of cancer-related death among men in the United States. In 2026, it is estimated that 333,830 men will be newly diagnosed with PC, while 36,320 will succumb to the disease [1]. When PC is confined to the prostate or regional lymph nodes, the 5-year survival rate is about 100% [2]. However, as the disease spreads to distant, metastatic sites, the 5-year survival rate drops to about 36% [2,3]. Multiple risk factors contribute to PC development, including age, ethnicity, obesity, and family history. Age is strongly linked to PC diagnosis, with the majority of newly diagnosed individuals over the age of 60 [4,5].

The prostate, an androgen-sensitive organ, depends on androgens and the androgen receptor (AR) signaling pathway for its growth. AR signaling is activated primarily by DHT, which is generated from testosterone by 5α-reductase and binds the androgen receptor (AR) with high affinity [6–8]. DHT then binds to the AR, leading to AR dimerization and phosphorylation [6,7,9]. Once phosphorylated, the receptor translocates to the nucleus to bind specific DNA sequences called androgen response elements (AREs) [6,7,10]. Here, AR acts as a transcription factor that regulates the expression of genes involved in cellular homeostasis and normal prostate function, including *KLK3*, which encodes prostate-specific antigen (PSA) [6]. Critically, dysregulated AR signaling pathways are a central driver of disease progression in benign prostatic hyperplasia (BPH) and PC. Tissue studies have confirmed AR expression across most PC subtypes, with AR detected in both epithelial cancer cells and the surrounding stromal compartment [11]. This broad distribution highlights the potential of AR signaling to regulate not only tumor intrinsic programs but also the tumor microenvironment.

In 1941, Huggins and Hodges established androgen deprivation as an effective therapeutic strategy for PC, demonstrating that PC growth could be substantially suppressed by reducing androgen signaling. Since then, androgen deprivation therapy, or ADT, has become a standard treatment for PC and remains a primary component of systemic therapy for advanced disease [7,12,13]. In its early years, ADT was mainly achieved through orchiectomy (surgical removal of the testicles), with pharmacological approaches emerging in the 1960s, including anti-androgens and estrogen therapy [7,14]. Later, in the 1980s, LHRH (also known as gonadotropin-releasing hormone, GnRH) agonists emerged as a non-surgical form of ADT that suppresses testosterone production by acting on the hypothalamic-pituitary-gonadal (HPG) axis [14–16]. These reversible forms of chemical castration have demonstrated overall survival rates similar to orchiectomy and, by 2009, were recommended by the National Comprehensive Cancer Network as first-line therapy for advanced disease [17–19]. However, GnRH agonists first stimulate pituitary GnRH receptors before desensitizing them. This causes an initial testosterone surge that can temporarily worsen symptoms and may stimulate tumor growth, particularly in patients with advanced disease [7,14,18,20].

GnRH antagonists were developed to avoid this flare [21]. Instead of first stimulating the pituitary GnRH receptor, antagonists block it directly, immediately suppressing luteinizing hormone (LH), follicle-stimulating hormone (FSH), and testosterone [14,22,23]. The GnRH antagonist Degarelix (FIRMAGON) results in rapid and sustained suppression of FSH, LH, and testosterone. It also lowers prostate-specific antigen (PSA) levels, a key biomarker used in PC diagnosis and surveillance [22]. In the clinic, Degarelix is given subcutaneously every 28 days, with a starting dose of 120 mg (twice) to immediately suppress testosterone, followed by smaller doses of 80 mg to keep the testosterone levels low [24]. ADT also has well-documented side effects beyond the prostate, including bone loss and increased fracture risk [25], metabolic changes such as insulin resistance and increased risk of diabetes [26], and loss of lean muscle mass with gains in fat mass [27,28].

Most patients initially respond to ADT with substantial clinical regression, reflected by falling PSA levels [3,29–31]. Despite this initial success, PSA eventually rises again as the disease progresses to castration-resistant prostate cancer or CRPC. This progression is driven largely by genetic alterations, many of which ADT itself induces or selects for. PC is genetically heterogeneous, with an accumulation of alterations, often copy number changes that vary between patients. Common alterations include deletions of *PTEN* and *RB1*, mutations in *TP53*, and amplifications of *MYC* and *AR* [32]. These alterations become enriched as the disease progresses. Amplifications and gain-of-function mutations in *AR* or amplifications of *AR* transcriptional regulators are present in about 70% of CRPC patients [33]. ADT frequently induces these alterations and can contribute to therapy resistance. Aberrations in genes involved in growth control also contribute to the progression of PC towards CRPC [3]. For example, while homozygous deletions of *PTEN* are only present in about 12-17% of localized PC, they are found in over 40% of CRPC tumors [33]. Furthermore, gain-of-function mutations in PI3K pathway intermediates have been shown to facilitate CRPC progression, and activation of the WNT signaling pathway is increased in CRPC [33–35]. Lastly, *TP53* and *RB1*, which control cell cycle arrest, are enriched in CRPC, present in 50% and 21%, respectively [33,34]. Consistent with clinical observations, studies in mouse models have shown that combined loss of *Pten, Trp53,* and *Rb1* promotes metastatic progression and lineage plasticity, supporting a role for these alterations in the development of aggressive CRPC [36].

Despite this, experimental models of CRPC are few and far between, with the most common CRPC models being cell lines derived from patients with CRPC. However, many of these models are effective only in immunocompromised animals, and many CRPC cell line models fail to recapitulate the true heterogeneity of the disease or represent treatment sensitivity [37,38]. Additionally, although genetically engineered mouse models are available, they can be labor-intensive and costly to generate and maintain, and often do not fully recapitulate the genetic and clinical complexity of CRPC [38]. There has been a recent push for 3D culture models, with success found with ex vivo tumor slice and organoid cultures [38]. However, these models are still in vitro systems and cannot model pharmacokinetics. Therefore, experimental immunocompetent models are still needed to study treatment response and resistance.

Developing such models requires a castration method that is both clinically relevant and well characterized in mice. Previous animal studies have demonstrated a castration-regeneration cycle where androgen withdrawal induced prostate involution and tissue regression, followed by regeneration and restoration of prostate size upon testosterone replacement [39]. Degarelix offers a clinically used, non-surgical option to withdraw androgens. However, the spatiotemporal dynamics of genitourinary tissue involution and regeneration after Degarelix induced androgen deprivation remain incompletely defined. The present study therefore used Degarelix-induced chemical castration to recapitulate the prostate involution-regeneration cycle. Utilizing immunocompetent mouse models, we evaluated the systemic effects of Degarelix and assessed its efficacy in reducing tumor burden.

## Materials and Methods

### Animal Studies

Male C57BL/6J mice (The Jackson Laboratory, Strain #000664) were maintained in our breeding colony. The UNMC IACUC approved all procedures. Degarelix Trial 1: Sixteen-week-old mice were randomized to continuous Degarelix (10 mg/kg, s.c., every 4 weeks), Degarelix withdrawal (single 10 mg/kg dose), or vehicle (n = 6/group). Blood was collected every 2 weeks by tail-snip bleed for serum testosterone measurement. Mice were euthanized at 4, 8, or 12 weeks, and urogenital tissues were collected for histological analysis. Degarelix Trial 2: Twelve-week-old mice received Degarelix (10 mg/kg, s.c.; n = 5) or vehicle (n = 5). Four days later, 2.5 x 10^5^ RapidCaP-derived *Pten/Trp53* ^NULL^ cells in Matrigel were injected subcutaneously. Tumor volume was measured weekly, and mice were euthanized at 4 weeks for tumor collection. Degarelix Trial 3: Twelve-week-old mice received 2.5 x 10^5^ luciferase-expressing RapidCaP-derived *Pten/Trp53* ^NULL^ cells in Matrigel by intraprostatic injection. Mice received Degarelix (10 mg/kg, s.c.; n = 5) or vehicle (n = 4) 12 days later. Tumor burden was monitored weekly by bioluminescence imaging, and mice were euthanized on day 47 for tissue collection. Intraprostatic injections were performed under isoflurane anesthesia using aseptic technique with perioperative buprenorphine (1 mg/kg, s.c.). Mice were monitored postoperatively until recovery and for 3 days thereafter.

### Generation of RapidCaP GEMM Cell Lines

Cell line models from the RapidCaP GEMM (*Pten/Trp53 ^NULL^*) were established as previously described in Naguib et al. [40]. Primary cell lines from RapidCaP mice were established through collagenase digestion of the prostate tissue. TdTomato-positive *Pten/Trp53 ^NULL^* cells were then isolated using fluorescence-activated cell sorting (FACS) [40]. These models have been subsequently utilized to study PC progression and therapeutic resistance [40–45].

### Cell Culture

Cells were maintained in RPMI medium supplemented with 2% fetal bovine serum (FBS), 100 µg/mL penicillin, 100 µg/mL streptomycin, and 2 mM L-glutamine. Cell lines were cultured as 2D monolayers in cell culture-grade Petri dishes and maintained at 37°C in a humidified incubator with 5% CO_2_. Routine checks for potential mycoplasma infection were conducted using the LookOut One-Step Mycoplasma PCR Detection Kit (Cat# MP0050-100TST) from Sigma-Aldrich.

### ELISA

Testosterone levels in mouse serum were measured using a Mouse Testosterone ELISA Kit (Crystal Chem, Cat# 80552) according to the manufacturer’s instructions. Serum samples were collected via tail-snip bleed, allowed to clot at room temperature, and centrifuged at 2000 g for 10 minutes at 4°C to separate the serum. Collected serum was stored at −80°C until ready to use. All reagents from the kit were brought to room temperature before use. First, 10 µL of sample, standard, or control was added to each well of a 96-well plate pre-coated with testosterone-specific antibodies. Next, 100 µL of Incubation Buffer was added to each well, followed by 50 µL of Enzyme Conjugate. The plate was then incubated for 1 hour at room temperature on a microplate shaker. After incubation, the plate was aspirated and washed 4 times with 300 µL of Wash Buffer (diluted 1:10 with deionized (DI) water) per well. Next, 200 µL of Substrate Solution was added to each well, and the plate was incubated for 30 minutes at room temperature in a dark room. The reaction was stopped by adding 50 µL of Stop Solution. The absorbance was then measured using a microplate reader (A_450_ – A_630_ values). A standard curve was generated using testosterone standard concentrations, and sample concentrations were calculated and plotted accordingly.

### Masson’s Trichrome staining

For Masson’s Trichrome staining, sectioned tissue slides were initially baked in a paraffin oven for 40 minutes at 62°C. They were then deparaffinized by first immersing them in three 5-minute changes of HistoChoice Clearing Agent (VWR Cat #97060-934), followed by two 5-minute changes in 100% ethanol, two in 95% ethanol, and one in 70% ethanol. The slides were then placed in DI water for 5 minutes. After deparaffinization and hydration, slides were stained with a Masson’s Trichrome Kit (Abcam Cat #ab150686) according to the manufacturer’s instructions. Whole slide scans of stained sections were obtained from the UNMC Tissue Sciences Facility. All images were acquired with identical exposure and white balance settings. Stromal thickness was quantified using ImageJ (version 1.54, National Institutes of Health, USA). Images were spatially calibrated using an embedded scale bar to convert pixels to micrometers. Stroma was identified as the collagen-rich, blue-stained region. Using the straight-line tool, thickness was measured perpendicular to the epithelial surface at 10 random areas per treatment condition.

### Immunofluorescence

Sectioned tissue slides were used for immunofluorescence staining. Slides were first baked in a paraffin oven for 40 minutes at 62°C. Slides were then deparaffinized, beginning with three 5-minute changes of HistoChoice Clearing Agent (VWR Cat #97060-934), followed by two changes of 100% ethanol, two changes of 95% ethanol, and one change of 70% ethanol, each for 5 minutes. Slides then underwent heat-induced antigen retrieval using either 10 mM sodium citrate (pH 6.0) or Tris-EDTA (pH 9.0), microwaved for 5 minutes at power 9, then 10 minutes at power 2. Slides were left to cool to room temperature before being moved to a moisture chamber. Slides were washed with PBS for 5 minutes, and boundaries were drawn around the tissue using a Dako Pen. Next, blocking buffer (10% normal goat serum (NGS) in PBS supplemented with 0.3% Triton X-100) was added to the slides for a 1-hour incubation. After 1 hour, the blocking buffer was removed and replaced with the primary antibody (Alexa Fluor 647 CK5, Abcam Cat# ab193895 1:200; Alexa Fluor 488 CK8, Abcam Cat# ab192467 1:200; NKX3.1, Athena Enzyme Systems Cat# 0314 1:50) diluted in 10% NGS in PBS with 0.3% Triton X-100. Slides were incubated with the primary antibody overnight at room temperature in a moisture chamber. The next day, the primary antibody was removed, and slides were washed 3 times with 1X PBS for 5 minutes each. For NKX3.1 staining, slides were then incubated with the appropriate fluorescently conjugated secondary antibody (Alexa Fluor 488; Invitrogen, Cat# A11008) for 1 hour at room temperature. The secondary antibody was then removed, and slides were washed with PBS 3 times for 5 minutes each. For all stains, slides were incubated with the nuclear stain DAPI (Sigma, Cat# D9542), diluted in DI water, for 10 minutes. After DAPI incubation, slides were washed with DI water and cover slipped with ProLong Gold mounting medium (Cell Signaling Technology, Cat# 9071S). Images of stained slides were acquired using the BioTek Cytation C10 Confocal Imaging Reader, with matched exposure and gain settings across groups for each stain. Images were taken at both 20x and 60x magnification (n = 10 random sections per tissue per stain; n = 1 mouse per group at week 4, n = 2 mice per group at week 8).

### Quantitative immunofluorescence analysis

Prostate sections collected at 4 and 8 weeks were stained for CK5, CK8, and DAPI, or NKX3.1 and DAPI, and imaged by confocal microscopy using a 60x objective. At 4 weeks, 40 CK5/CK8 fields and 41 NKX3.1 fields were analyzed from vehicle- and Degarelix-treated animals. At 8 weeks, 61 fields were analyzed for each marker set across vehicle, castration, and regeneration groups. The field of view (∼210 µm) was the unit of analysis. Nuclei were segmented from DAPI images using adaptive thresholding and watershed separation. For CK5/CK8 analyses, epithelial nuclei were identified from the combined CK5/CK8 signal; all nuclei were analyzed for NKX3.1. CK5 and CK8 fluorescence was measured within a 2.4-µm perinuclear annulus, whereas NKX3.1 was measured within the nuclear mask and normalized to local tissue signal. CK5 positivity was defined using the field-specific Otsu threshold, CK8 positivity using a fixed intensity threshold of 10 grey levels, and NKX3.1 positivity as nuclear signal ≥1.1-fold the local tissue reference across ≥60% of the nucleus. H-scores (0–300) were calculated as 1 × %weak + 2 × %moderate + 3 × %strong. Intensity grades were based on within-field normalized measurements, with cut-points anchored to the corresponding vehicle cohort at each timepoint. Analyses were performed using Python 3.13 with scikit-image, SciPy, NumPy, and pandas. Image segmentation, marker quantification, and H-score analysis were performed using Claude (Anthropic, San Francisco, CA), an AI-assisted research tool.

## Statistical Analysis

All statistical analyses were performed using GraphPad Prism 10.4.1 (GraphPad Software, Inc.; Boston, MA). Comparison of stromal thickness between vehicle, castration, and regeneration cohorts was evaluated using one-way ANOVA with Dunnett’s multiple comparisons. For immunofluorescence data, two-group comparisons at 4 weeks used two-sided Mann-Whitney U tests. Three-group comparisons at 8 weeks used Kruskal-Wallis tests followed by Holm-adjusted pairwise Mann-Whitney U tests. Effect size was reported as AUC. Comparisons of mouse serum testosterone levels, body weight, and tumor growth response between treatment conditions were evaluated using two-way ANOVA with Šidák’s multiple comparisons test. Comparison of tumor weight ex-vivo between treatment conditions was evaluated using a Mann-Whitney U test. All results were reported as mean ± SEM. For all tests, p-values < 0.05 were considered statistically significant.

## Results

To evaluate the physiological effects of Degarelix induced chemical castration *in vivo*, we randomized C57BL/6J mice into three groups: a control group (vehicle-treated), a short-term Degarelix treatment group (Deg^ST^), and a long-term Degarelix treatment group (Deg^LT^). Animals in the Deg^ST^ cohort received a single dose of Degarelix (10 mg/kg) for 4 weeks, followed by an 8-week drug-free period to assess recovery of androgen suppression and testosterone levels (Fig. 1a). As shown in Fig. 1a, animals in the Deg^LT^ cohort received monthly Degarelix (10 mg/kg) for 12 weeks to assess sustained androgen deprivation. Animals were sacrificed at the indicated time points, as shown in the schematic in Fig. 1a.

**Figure 1:**
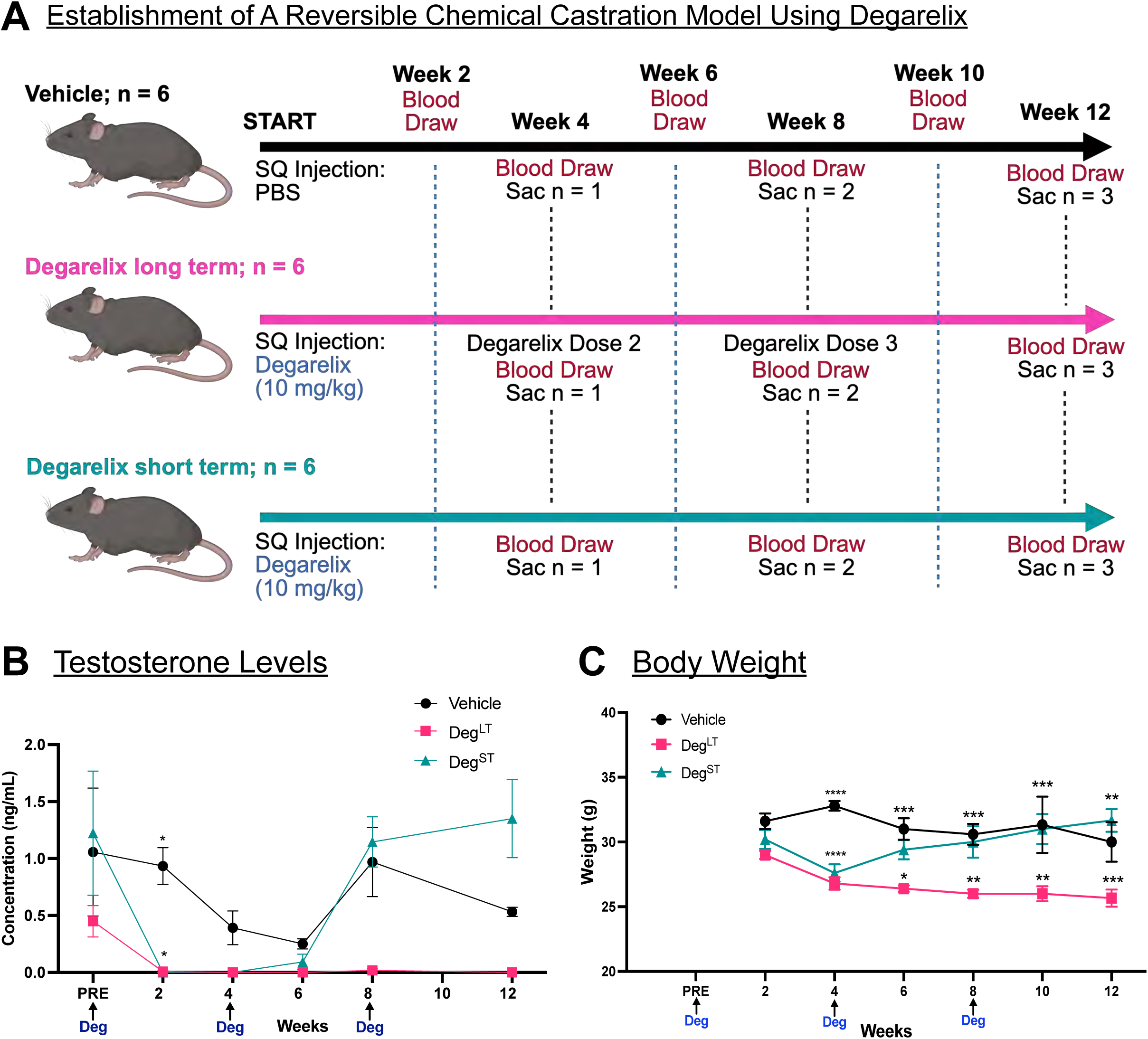
Prolonged treatment with Degarelix significantly reduced testosterone levels and body weight, whereas Degarelix removal restored both parameters, demonstrating the reversibility of chemical castration. **(A)** Experimental strategy for evaluating the systemic effects of Degarelix *in vivo*. **(B)** A significant reduction in testosterone levels was observed following the initial dose of Degarelix, with levels remaining suppressed (near 0.000 ng/mL) during long-term treatment and increasing following Degarelix withdrawal (short-term Degarelix); n=3 animals per group at each time point. **(C)** Long-term Degarelix treatment causes significant weight loss. Mice receiving short-term Degarelix showed significant initial weight loss, which was reversed following treatment withdrawal. n=6 animals per group for weeks 2-4; n=5 animals per group for weeks 5-8; n=3 animals per group for weeks 9-12. **** represents p<0.0001; *** represents p<0.001; ** represents p<0.01; * represents p<0.05; two-way ANOVA with Šidák’s multiple comparisons. Deg^ST^ refers to the Degarelix short-term cohort, and Deg^LT^ refers to the Degarelix long-term cohort.

As a GnRH antagonist, Degarelix rapidly suppresses luteinizing hormone and testosterone without the surge associated with GnRH agonists, providing a pharmacologic model of sustained androgen deprivation [46,47]. We therefore monitored circulating testosterone throughout the study to confirm the efficacy and durability of androgen suppression in the Deg^ST^ and Deg^LT^ cohorts. Blood samples were collected every 2 weeks for measurement of serum testosterone levels by ELISA. As early as two weeks post Degarelix treatment, serum testosterone levels decreased significantly in the Deg^LT^ and Deg^ST^ groups relative to vehicle controls (0.008 ± 0.008 ng/mL vs 0.934 ± 0.161 ng/mL; p < 0.05; Fig. 1b). Testosterone levels in the Deg^LT^ cohort remained near the lower limit of detection throughout the study, consistent with sustained suppression of systemic testosterone levels (Fig. 1b). In contrast, following Degarelix withdrawal, serum testosterone in the Deg^ST^ cohort rose to levels comparable to those in the vehicle group (1.145 ± 0.231 ng/mL vs 0.970 ± 0.304 ng/mL; Fig. 1b). These findings demonstrate that testosterone suppression induced by Degarelix is reversible following treatment withdrawal.

Androgen deprivation is associated with systemic metabolic changes that can affect body-weight trajectories [27,48]. We therefore assessed body weight following Degarelix treatment. One-month post Degarelix treatment, we observed a significant decrease in body weight in the Deg^LT^ and Deg^ST^ cohorts compared with the vehicle cohort (26.800 ± 0.490 g and 27.600 ± 0.678 g vs 32.800 ± 0.374 g; p < 0.0001 for both; Fig. 1c). Following Degarelix withdrawal, body weight in the Deg^ST^ cohort began to recover by 2 weeks and, by 8 weeks, and was similar to that of vehicle-treated animals (30.000 ± 1.225 g vs 30.600 ± 0.812 g; Fig. 1c). In contrast, the initial decline in body weight in the Deg^LT^ cohort was sustained throughout the trial. At the terminal endpoint (12 weeks), body weight in the Deg^LT^ cohort remained significantly lower than that in both the vehicle and Deg^ST^ cohorts (Fig. 1c). Thus, unlike the weight gain commonly reported in PC patients receiving androgen deprivation therapy [27,49,50], Degarelix-induced castration in healthy, disease-free C57BL/6J mice resulted in significant weight loss. This effect on body weight was reversible after treatment withdrawal but persisted with continued treatment. Because the association between androgen deprivation and weight gain has been established primarily in men with PC, these findings highlight the need to better define the metabolic effects of androgen deprivation in preclinical models of PC.

Next, to assess the effects of Degarelix on prostate morphology, we performed histological analysis on the urogenital system collected from mice in each cohort at weeks 4, 8, and 12. The urogenital system, including the bladder and intact anterior, ventral, and dorsolateral lobes of the prostate, was dissected and processed for histopathology. At the 8-week time point, H&E-based histological analysis demonstrated significant shrinkage of the prostate following Degarelix treatment and restoration of prostate size after Degarelix withdrawal, consistent with the reversible effects of pharmacological androgen deprivation (Supplementary Fig. 1a-c). Consistent with these findings, Masson’s trichrome staining of prostate tissue from animals in the vehicle cohort (8-week) showed normal prostate histopathology, with stromal tissue exhibiting the characteristic blue staining (Supplementary Fig. 2a). In contrast, Masson’s Trichrome staining of Degarelix-treated prostate tissue revealed an expanded stromal layer following castration compared with vehicle-treated controls (0.635 ± 0.075 µm vs 0.249 ± 0.308 µm; Supplementary Fig. 2b, d). Following Degarelix withdrawal and prostate regeneration, stromal layer thickness decreased to 0.272 ± 0.033 µm, similar to prostates from animals in the vehicle-treated cohort (Supplementary Fig. 2c, d). Collectively, our data show that Degarelix treatment led to prostate involution with stromal layer thickening, and Degarelix withdrawal restored normal prostate size and stromal architecture.

To determine how Degarelix induced androgen deprivation and its withdrawal affect the prostate epithelium, we performed immunofluorescence analysis of basal and luminal lineage markers. We also assessed *NKX3.1*, a prostate-specific homeobox gene whose expression is androgen-dependent [51], as a readout of AR signaling. The prostate epithelium consists of two principal lineages: CK5-positive basal cells and CK8-positive luminal (glandular) cells. Androgen deprivation affects the two cell types differently; castration preferentially eliminates androgen-dependent luminal cells, whereas basal cells are largely spared and persist in the regressed gland [52,53]. In addition, androgens regulate cytokeratin expression in the prostate. Following castration, expression of the luminal cytokeratins CK8 and CK18 and the basal cytokeratins CK5 and CK15 increased in the rat ventral prostate, identifying these genes as a class of androgen-repressed genes; CK8 immunostaining similarly shifted from a uniform luminal distribution to predominantly basal and transit-amplifying cells [54]. Consistent with these reports, immunofluorescence staining for the basal and luminal markers CK5 and CK8 showed increased CK5-positive cells and more robust CK8 staining following Degarelix-induced castration at 4 weeks (Fig. 2a). In contrast, staining for the AR target gene *NKX3.1* showed reduced nuclear expression at 4 weeks, confirming effective suppression of AR signaling by Degarelix (Fig. 2b). The fraction of CK5-positive nuclei within the epithelium increased 2.01-fold, from 13.27% to 26.65% following castration (p = 4.7×10^-5^; Fig. 2c), while the CK5 H-score increased 1.43-fold, from 26.25 to 37.51 (p = 3.4×10^-4^, Fig. 2d). The CK8 H-score was unchanged between cohorts, with values of 198.63 in vehicle-treated animals and 200.00 following castration (1.01-fold, p = 0.34; Fig. 2e). CK8 was detected in 100% of epithelial nuclei in both groups, indicating that total keratin coverage was preserved despite the shift in CK5 abundance. The H-score for nuclear NKX3.1 decreased by 60%, from 108.70 in vehicle-treated animals to 43.32 following castration (0.399-fold, p = 1.2×10^-5^; Fig. 2f).

**Figure 2:**
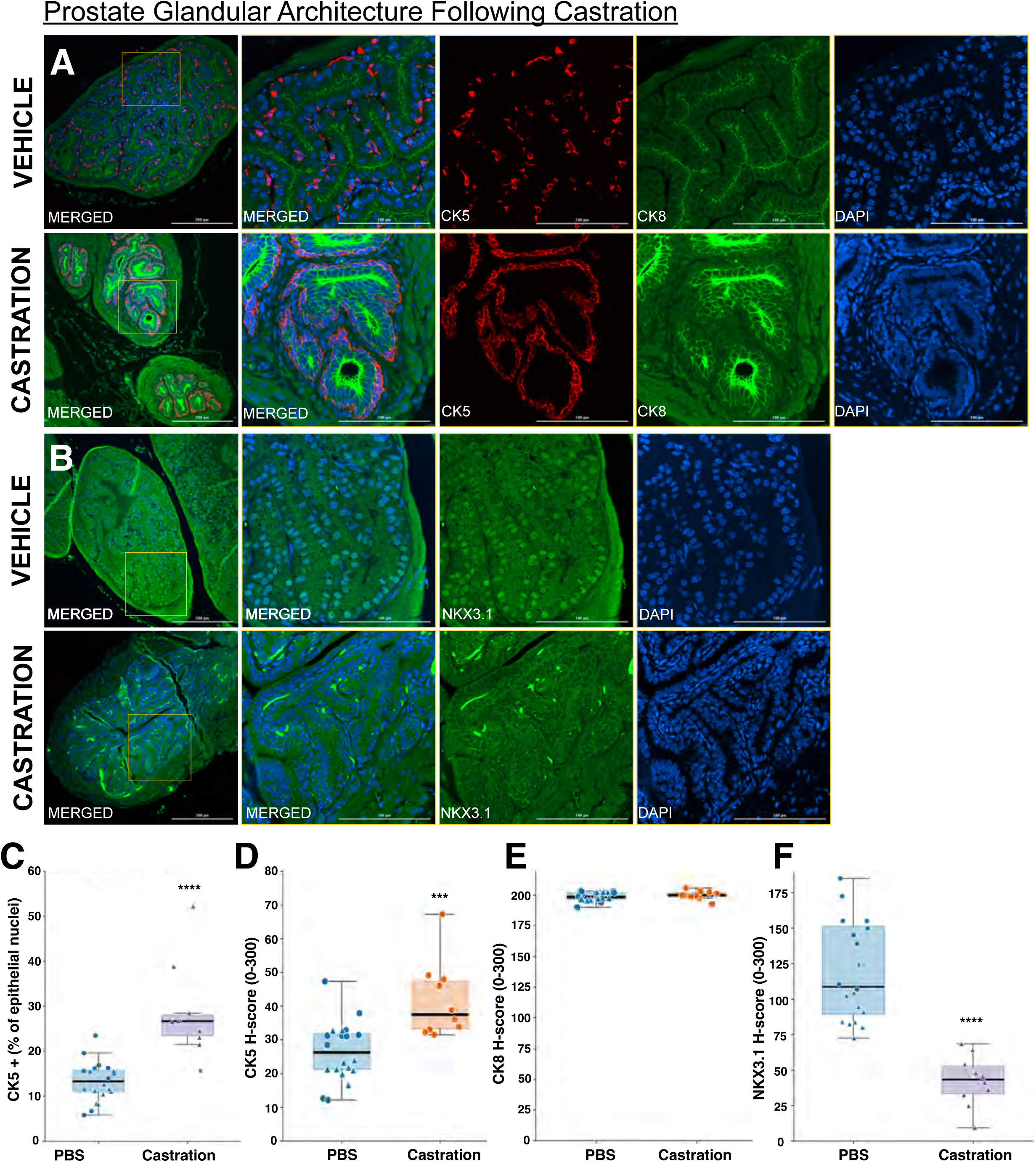
Degarelix-induced castration alters CK5-positive basal cell populations and reduces NKX3.1 expression in the prostate epithelium. **(A)** Degarelix-induced castration qualitatively increases CK5-positive cells and enhances CK8 staining in the prostate epithelium compared with vehicle-treated tissue. **(B)** IF staining of vehicle-treated prostate tissue showed normal androgen receptor activity, indicated by abundant NKX3.1-positive cells. Following Degarelix treatment, NKX3.1 expression within the prostate was markedly reduced, consistent with suppression of androgen receptor signaling. Representative images of each group at the 4-week time point. Leftmost merged images: Magnification = 20x, scale bar = 200 µm. The images to the right are zoomed-in views of the areas indicated by the yellow squares; magnification = 60x, scale bar = 100µm. Images were acquired using the BioTek Cytation C10 Confocal Imaging Reader, with matched exposure and gain settings across groups for each stain. **(C)** The CK5-positive fraction of the prostate epithelium doubles after castration. Quantification of CK5-positive nuclei as a percentage of nuclei within the prostatic epithelium; 60x magnification. Vehicle: n=20 sections from 2 animals; castration: n=10 sections from 1 animal. **(D)** Castration increases CK5 expression in the prostate epithelium. **(E)** CK8 expression was detected in the transit-amplifying cell population in prostate tissue from castrated animals. **(F)** Castration reduces nuclear NKX3.1 expression in the prostate epithelium. H-score was calculated per field; vehicle: n=20 sections from 2 animals, castration: n=10 sections from 1 animal. Boxes indicate the interquartile range with the median shown; whiskers extend to 1.5xIQR. p-values reflect section-level variation. **** represents p < 0.0001, *** represents p < 0.001; two-sided Mann-Whitney U test.

At 8 weeks, CK5 and CK8 immunofluorescence showed that, following prostate regeneration, the abundance and distribution of CK5- and CK8-positive cells in the Deg^ST^ cohort were comparable to vehicle-treated animals (Fig. 3a-b). Similarly, following Degarelix withdrawal and prostate regeneration, NKX3.1 staining showed restoration of NKX3.1-positive nuclei within the epithelium to levels comparable to vehicle-treated animals, indicating recovery of androgen signaling. (Fig. 3c-d). The fraction of CK5-positive nuclei within the epithelium increased 1.77-fold, from 20.83% in vehicle-treated animals to 36.78% following castration (p = 9.0×10^-4^; Fig. 3e). Following regeneration, CK5-positive nuclei decreased to 19.08%, comparable to vehicle-treated animals (p = 0.49) and significantly lower than after castration (p = 3.9×10^-4^; Fig. 3e). Consistent with these findings, the CK5 H-score increased 1.68-fold with castration, from 41.07 to 69.19 (p = 8.2×10^-4^), and returned to 38.99 following regeneration, comparable to vehicle-treated animals (p = 0.58; Fig. 3f). CK8 H-score was unchanged across vehicle, castration, and regeneration groups (200.36, 199.19, and 198.16, respectively; p = 0.19), with CK8 detected in 100% of epithelial nuclei across all groups (Fig. 3g). The NKX3.1 H-score decreased from 108.90 in vehicle-treated animals to 80.15 following castration (0.736-fold, p = 9.6×10^-5^) and increased to 155.81 after regeneration (1.43-fold vs. vehicle, p = 1.3×10^-6^; Fig. 3h). The fraction of NKX3.1-positive nuclei similarly decreased from 55.2% to 44.4% with castration and increased to 66.2% after regeneration. Together, these data show that Degarelix-induced androgen deprivation suppresses AR signaling and shifts the prostate epithelium toward a relatively basal-enriched state. Both epithelial composition and NKX3.1 expression recover following Degarelix withdrawal. This pattern is consistent with the castration/regeneration cycle described after surgical castration and androgen replacement [53,55]. Thus, Degarelix treatment and withdrawal provide a reversible, non-surgical model of androgen deprivation and prostate regeneration.

**Figure 3:**
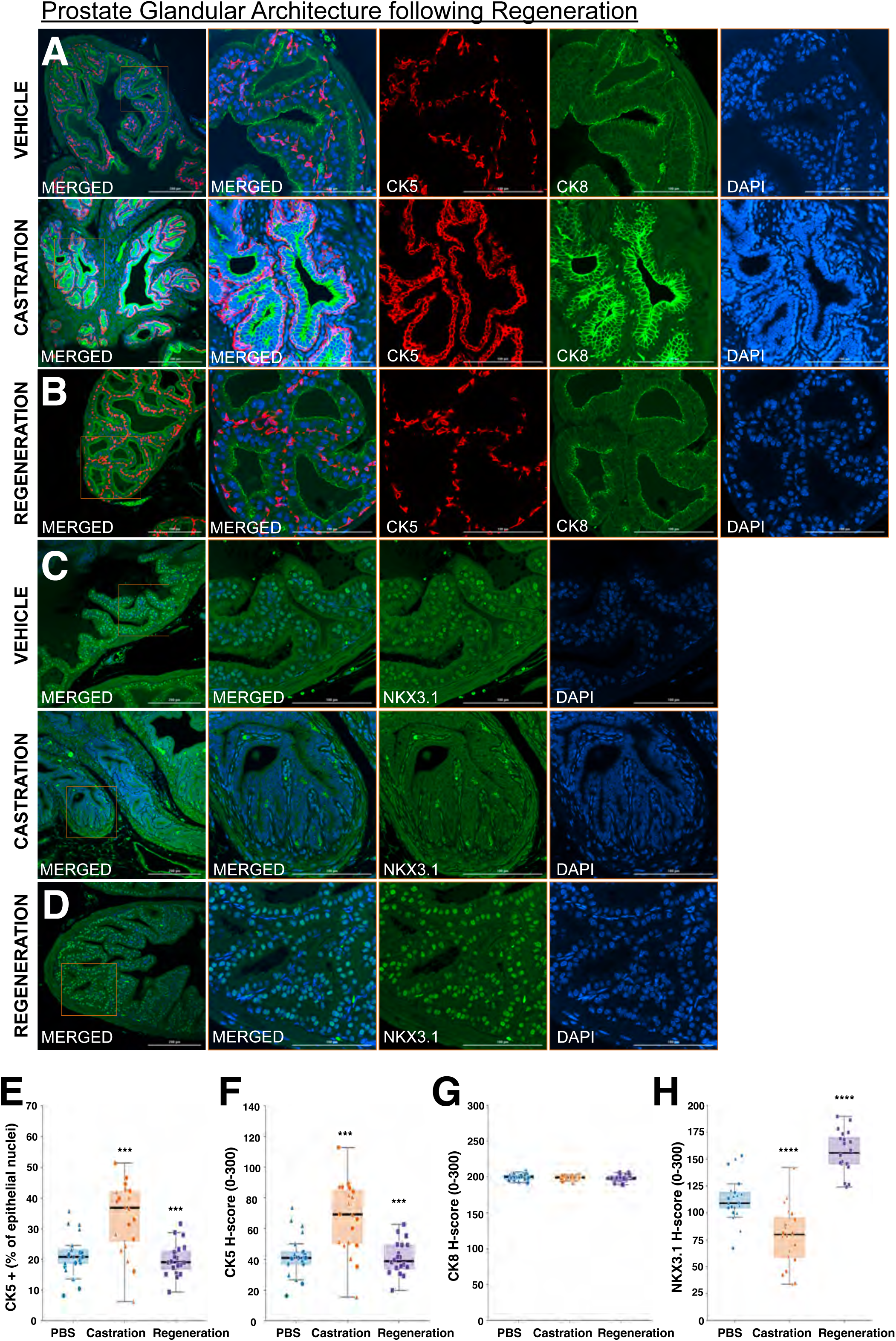
Degarelix withdrawal restores CK5- and CK8-positive staining and NKX3.1 expression in the prostate epithelium. **(A)** Prolonged Degarelix treatment (castration) increases CK5-positive basal cell populations within the prostate epithelium compared with vehicle-treated tissue. **(B)** Four weeks after Degarelix withdrawal (regeneration), CK5- and CK8-positive staining returned to levels comparable to vehicle-treated tissue. **(C)** IF staining of vehicle-treated tissue showed intact androgen receptor signaling, indicated by abundant NKX3.1-positive cells. Following Degarelix treatment (castration), NKX3.1-positive staining was markedly reduced, consistent with suppressed androgen receptor signaling. **(D)** Staining of prostate tissue 4 weeks after Degarelix withdrawal (regeneration) showed restoration of NKX3.1-positive cells, consistent with restored androgen receptor signaling. Representative images from each group are shown at the 8-week time point. Leftmost merged images: Magnification = 20x, scale bar = 200 µm. The images to the right are zoomed-in views of the areas indicated by the orange squares; magnification = 60x, scale bar = 100µm. Images were acquired using the BioTek Cytation C10 Confocal Imaging Reader, with matched exposure and gain settings across groups for each stain. **(E)** Castration expands the CK5-positive epithelial population, and regeneration reverses it. Quantification of CK5-positive nuclei as a percentage of nuclei within the prostatic epithelium; 60x magnification. Vehicle: n=20 sections from 2 animals; castration: n=20 sections from 2 animals; and regeneration: n=20 sections from 2 animals. **(F)** Castration increases CK5 expression, and regeneration reverses it. **(G)** CK8 expression was preserved across all three cohorts, with CK8-positive cells observed in the transit-amplifying population following castration. **(H)** Nuclear NKX3.1 expression decreased with castration and was restored following regeneration. H-score was calculated per field; v Vehicle: n=20 sections from 2 animals; castration: n=20 sections from 2 animals; and regeneration: n=20 sections from 2 animals. Boxes indicate the interquartile range with the median shown; whiskers extend to 1.5xIQR. p-values reflect section-level variation. **** represents p < 0.0001, *** represents p < 0.001; two-sided Mann-Whitney U test.

Having established that Degarelix induces androgen deprivation and that this effect is reversible, we next evaluated Degarelix-induced ADT in preclinical models of PC that more closely recapitulate the clinical course of ADT. *PTEN* and *TP53* are among the most frequently inactivated tumor suppressors in human PC [56,57], and their combined loss defines an aggressive, therapy-resistant subset of the disease, making this genotype a clinically relevant context in which to model ADT response. RapidCaP is a genetically engineered mouse model (GEMM) in which *Pten* and *Trp53* are conditionally deleted in the adult prostate through viral delivery of Cre recombinase [58]. In the *Pten* null setting, loss of *Trp53* promotes distant metastasis in more than 50% of mice within 4 months [58]. Importantly, RapidCaP recapitulates key features of human PC, including an initial response to castration followed by the development of castration-resistant disease associated with a switch from AKT to MYC signaling [58]. We isolated cell lines from RapidCaP tumors [40] and established an immunocompetent subcutaneous experimental model. In a pilot trial, *Pten/Trp53* ^NULL^ cells were implanted subcutaneously to define tumor growth kinetics and establish the initial response to Degarelix induced ADT. C57BL/6J mice were randomized into two cohorts: a Degarelix-pretreated cohort (10 mg/kg) and a vehicle-pretreated control cohort (Fig. 4a). Four days after administration of Degarelix or vehicle (PBS), *Pten/Trp53* ^NULL^ cells were injected subcutaneously into the right flank of each animal (Fig. 4a). We monitored tumor burden weekly by caliper measurements and collected blood weekly to assess serum testosterone levels. Consistent with our findings in healthy mice, testosterone levels were substantially reduced in the Degarelix-treated cohort compared with the vehicle control group. Testosterone levels declined following Degarelix administration and remained suppressed throughout the trial (Fig. 4b). Degarelix treatment initially inhibited tumor growth in *Pten/Trp53* ^NULL^ cells, with a significant reduction in tumor volume observed at 3 weeks post-implantation (Fig. 4c). However, by the 4-week terminal endpoint, tumor volumes in the Degarelix-treated and vehicle control groups were comparable, consistent with reduced sensitivity to androgen deprivation and potentially reflecting the development of acquired resistance (Fig. 4c). As in disease-free mice, body weight was significantly reduced with Degarelix treatment at weeks 3 and 4; however, tumor weights of tumors excised at the trial endpoint did not differ between the Degarelix- and vehicle-treated groups (Fig. 4d, e).

**Figure 4:**
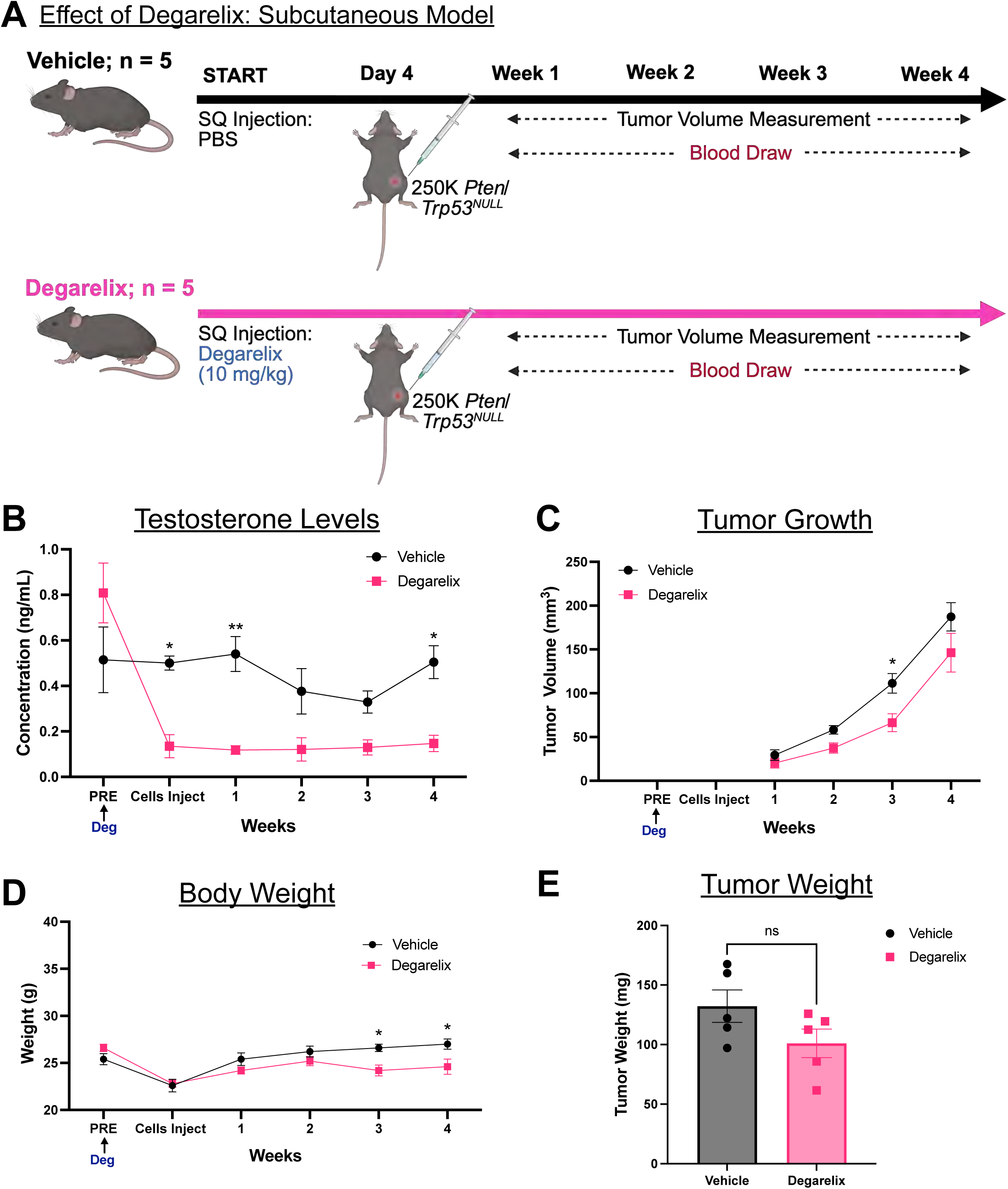
Degarelix-induced androgen deprivation initially inhibits tumor growth, followed by acquired resistance. **(A)** Experimental timeline for evaluating the effects of Degarelix on subcutaneous PC tumor growth. **(B)** Degarelix treatment significantly suppressed testosterone levels, which remained near 0.000 ng/mL throughout the study. n=4 animals for the vehicle group; n=3 animals for the Degarelix-treated group. ** represents p<0.01; * represents p<0.05; two-way ANOVA with Šidák’s multiple comparisons. **(C)** Degarelix-induced castration initially suppressed PC tumor growth, with a significant reduction at 3 weeks. By the trial endpoint, tumor growth was comparable to vehicle-treated controls, consistent with acquired resistance. * represents p<0.05; two-way ANOVA with Šidák’s multiple comparisons. **(D)** Degarelix treatment reduced body weight, with significant reductions at Weeks 3 and 4. * represents p<0.05; two-way ANOVA with Šidák’s multiple comparisons. **(E)** Ex vivo tumor weight did not differ significantly between treatment groups. n = 5 animals per cohort; ns, not significant; Mann–Whitney U.

Given the limitations of subcutaneous tumors, which lack the native prostate microenvironment, we next developed an immunocompetent orthotopic intraprostatic model to better recapitulate the disease context and assess the response to Degarelix and the development of treatment resistance. To longitudinally and noninvasively monitor tumor burden, we engineered the cells to express luciferase, enabling quantification of tumor growth by bioluminescence imaging (BLI). We injected luciferase-expressing *Pten/Trp53* ^NULL^ cells orthotopically into the anterior prostate of C57BL/6J mice (Fig. 5a). BLI of representative mice one day after surgery confirmed successful injection of tumor cells into the anterior prostate (Fig. 5b). On day 12, following confirmation of tumor establishment by BLI, mice were randomly divided into two groups, with half receiving Degarelix and the other half receiving PBS as the vehicle control (Fig. 5a-b). BLI revealed comparable tumor burden between the two cohorts at early time points following tumor implantation (Fig. 5c-d). A significant reduction in BLI signal was observed in the Degarelix-treated cohort on days 33 and 40, indicating an initial response to Degarelix-induced ADT (Fig. 5b, e). However, this difference was no longer evident by day 47, with BLI signal in the Degarelix-treated cohort similar to that of the vehicle-treated cohort, consistent with a diminished response to androgen deprivation and potentially the development of acquired resistance (Fig. 5c, f). To maintain androgen deprivation throughout the study, we administered a second dose of Degarelix on day 40, prior to the trial endpoint on day 47. Together, our data demonstrate that Degarelix induced ADT transiently suppresses tumor growth in the orthotopic *Pten/Trp53* ^NULL^ model, followed by tumor regrowth despite sustained testosterone suppression, consistent with the development of acquired resistance to androgen deprivation.

**Figure 5:**
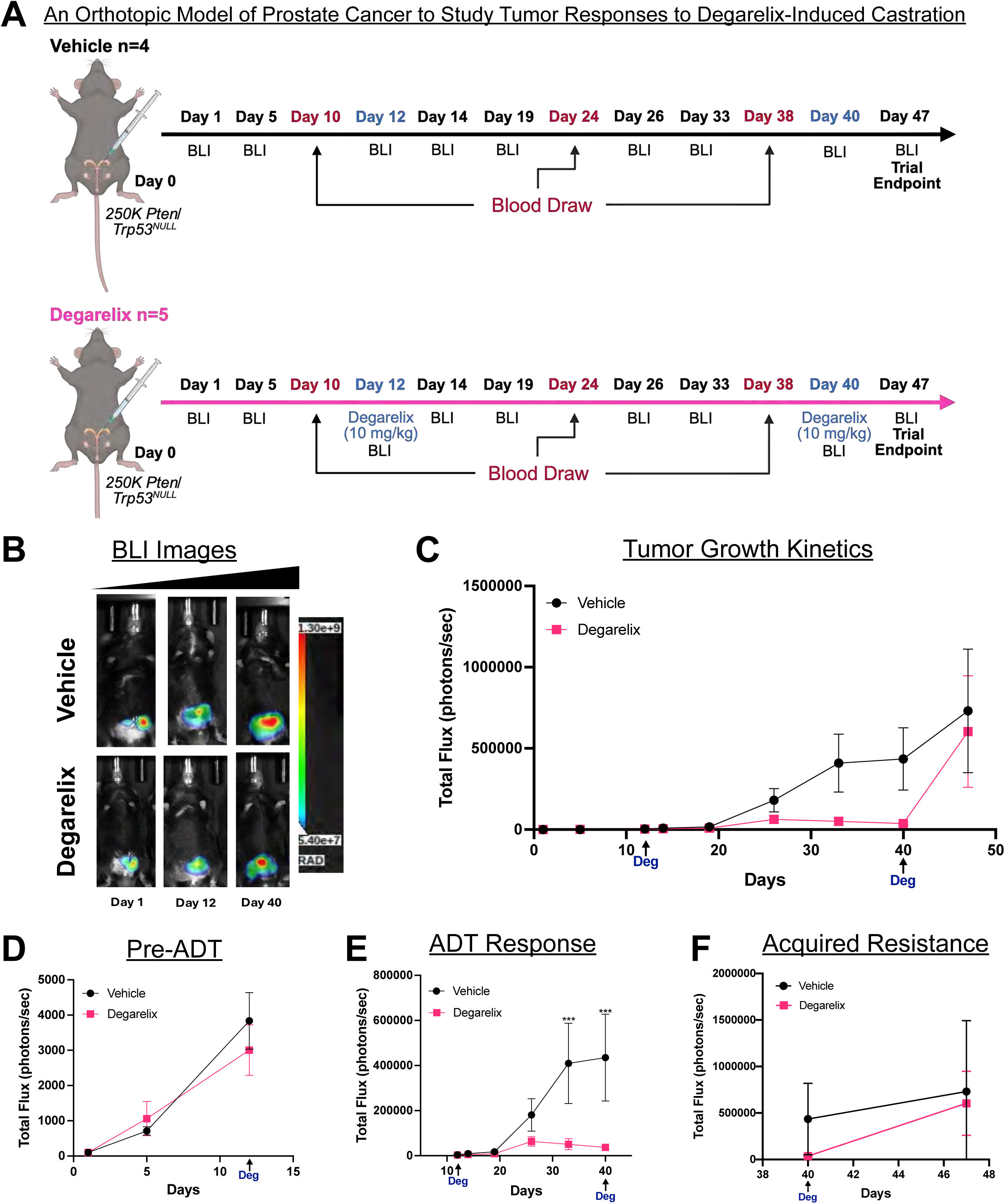
Orthotopic prostate cancer tumors develop resistance to Degarelix-induced castration. **(A)** Experimental timeline for evaluating the effects of Degarelix on orthotopic PC tumor growth. Bioluminescence imaging days are indicated in black; blood collection days in red; and Degarelix treatments in blue. **(B)** Representative bioluminescence images of vehicle- and Degarelix-treated mice one day after cell line transplantation and on Days 12 and 40 of Degarelix treatment. **(C)** Degarelix treatment initially reduced tumor growth; however, by the trial endpoint, tumor burden was comparable between vehicle and castration groups. Bioluminescence imaging (BLI) was used to longitudinally monitor tumor growth. **(D)** Before Degarelix treatment on Day 12, BLI showed no significant difference between cohorts. **(E)** Following Degarelix treatment, BLI signal in the Degarelix-treated cohort showed significantly reduced tumor growth on Days 33 and 40, indicating an initial treatment response. **(F)** By the trial endpoint (Day 47), BLI signal showed no significant difference in tumor burden between cohorts, consistent with acquired resistance to castration. n = 4 animals in the control group; n = 5 animals in the Degarelix-treated group. *** represents p>0.001; two-way ANOVA with Šidák’s multiple comparisons.

Collectively, these studies establish Degarelix induced chemical castration as a reversible, non-surgical model of androgen deprivation in mice and extend its application to preclinical models of PC. In healthy C57BL/6J mice, a single dose of Degarelix suppressed serum testosterone to near the limit of detection. We show that systemic androgen deprivation induced prostate gland involution, stromal expansion, loss of NKX3.1 expression, and a shift toward a basal cell-enriched epithelium. These changes reversed following drug withdrawal, whereas continuous dosing sustained androgen suppression and weight loss through 12 weeks. In immunocompetent preclinical models of PC generated using *Pten/Trp53* ^NULL^ tumors, Degarelix produced an initial anti-tumor response, with significant reductions in tumor volume and BLI signal at early time points. However, tumor growth resumed by the trial endpoint despite continued testosterone suppression, consistent with the development of acquired resistance to androgen deprivation. Together, these findings establish a physiologically relevant platform for modeling androgen deprivation and acquired resistance *in vivo*, providing a foundation for dissecting resistance mechanisms and evaluating therapeutic strategies to overcome treatment resistance.

## Discussion

PC remains the most commonly diagnosed cancer in men, with a lifetime risk of 13.2% [1]. Since the seminal observation that androgen withdrawal induces regression of metastatic PC, suppression of AR signaling through ADT has remained the backbone of treatment for both localized and advanced disease [12]. Despite an initial response in most patients, approximately 10-20% progress to castration-resistant prostate cancer (CRPC) within five years of diagnosis. Although treatment options for CRPC have expanded considerably, the disease remains incurable [59–61]. Defining the mechanisms that drive this transition requires preclinical models in which androgen deprivation can be precisely applied, sustained, and reversed in a controlled and physiologically relevant manner.

Degarelix, a GnRH antagonist, is a potent suppressor of the male hormones testosterone, FSH, and LH. FSH has been associated with PC growth and with the progression to CRPC [62]. Therefore, Degarelix offers an advantage over GnRH agonists, in which FSH levels rebound after initial suppression [62], a phenomenon ADT-resistant cancer cells can exploit to promote growth and survival. Degarelix provides a pharmacologic means of achieving sustained, single-agent suppression of the HPG axis, providing a rationale for characterizing its systemic effects *in vivo*.

In this study, we show that a single dose of Degarelix suppressed serum testosterone to near the limit of detection within 2 weeks, while monthly dosing maintained this suppression for 12 weeks. Sustained androgen deprivation induced prostate involution accompanied by stromal expansion, loss of nuclear NKX3.1, and a shift toward a relatively basal-enriched epithelial layer. These features recapitulate the classical response to surgical castration. Castration induced involution reflects the preferential loss of androgen-dependent luminal cells, whereas basal cells are relatively spared and persist in the regressed gland [52,53]. The increase in CK5-positive basal cells and the intense CK8 staining in the luminal compartment observed following Degarelix treatment are consistent with evidence that androgen withdrawal expands the basal and basal/luminal intermediate compartments of the prostatic epithelium [52,63,64]. Notably, CK8 redistributes from a predominantly luminal to a more basal pattern after androgen withdrawal [54]. Similarly, the loss and subsequent recovery of NKX3.1 staining are consistent with the established androgen dependence of this prostate-specific homeobox gene [51].

Importantly, we show that these changes were reversible following Degarelix withdrawal, with prostate glands regenerating to sizes comparable to those of vehicle-treated controls. These results are consistent with published literature showing ductal regeneration following testosterone propionate administration to castrated mice [55] and establish Degarelix treatment and withdrawal as a reversible, non-surgical model of the castration-regeneration cycle. The key conceptual advantage of this approach is how it restores androgen signaling. Conventional regeneration protocols rely on exogenous testosterone delivered by injection or subcutaneous pellet, which bypasses the HPG axis and can suppress GnRH, LH, and FSH, with downstream effects on spermatogenesis and fertility [65]. In contrast, Degarelix clearance permits recovery of endogenous testosterone production through the intact HPG axis, providing a more physiological model of androgen restoration. This distinction is particularly relevant to bipolar androgen therapy (BAT), in which cycling between supraphysiologic and castrate testosterone levels has been investigated as a strategy to resensitize CRPC tumors to subsequent androgen deprivation [66,67]. A model in which androgen levels can be pharmacologically lowered and subsequently restored without exogenous hormone administration therefore provides a tractable system for dissecting tumor responses to androgen withdrawal and re-exposure.

An unanticipated finding was the divergent effect of Degarelix treatment on body weight in mice. ADT in men is consistently associated with increased fat mass and weight gain [48,49], whereas Degarelix-treated healthy C57BL/6J mice exhibited significant weight loss. This effect was reversible following treatment withdrawal and persisted with continuous dosing. Several factors may contribute to this apparent species discordance. The mice used here were young adults (3-4 months of age), whereas men receiving ADT are typically several decades older and have age-related changes in body composition, including reduced lean mass and increased adiposity, that are not present to the same extent in young animals. Moreover, young mice remain on an anabolic growth trajectory; androgen deprivation may therefore attenuate ongoing lean mass accrual rather than produce net catabolic loss, resulting in a direction of body-weight change that differs from that observed in an older, weight-stable population. Other contributing factors may include species-specific metabolic responses to androgen deprivation, the absence of tumor burden and its associated systemic effects in our model, and the relatively short study duration compared with the years over which body-composition changes develop in patients. Because the metabolic consequences of ADT have been studied predominantly in men with PC, our findings underscore that changes in body weight following androgen deprivation may not translate directly between mice and humans. These findings also support systematic assessment of body composition, including fat mass, lean mass, and metabolic parameters, rather than body weight alone in preclinical studies of androgen deprivation and, where feasible, evaluation of aged animals to better model the metabolic context of the patient population.

We next evaluated the effects of Degarelix induced castration in tumor-bearing mice. *PTEN* and *TP53* are among the most frequently inactivated tumor suppressors in PC [56,57], and the RapidCaP model recapitulates key features of the human disease, including metastatic dissemination, an initial response to castration, and subsequent castration-resistant progression associated with a switch from AKT to MYC signaling [58]. We show that, in both subcutaneous and orthotopic models, Degarelix induced a significant early reduction in tumor burden that was no longer apparent at the study endpoint. Because serum testosterone remained suppressed throughout the treatment period, the subsequent tumor regrowth is unlikely to reflect loss of pharmacologic activity and instead reflects the emergence of acquired resistance to androgen deprivation. These findings extend prior studies in RapidCaP GEMM demonstrating the development of CRPC following surgical castration [58] and show that this trajectory can also be recapitulated through pharmacologic androgen deprivation. Importantly, both models are immunocompetent, providing an experimental context in which tumor-immune interactions can be incorporated into studies of ADT response and resistance.

This study has several limitations. Cohort sizes were modest, and although the effects of Degarelix on prostate remodeling were consistent across animals and tumor response patterns were reproduced in two independent model systems, larger studies are needed to confirm these findings and detect more subtle intergroup differences. Our resistance phenotype is currently defined by tumor growth kinetics under sustained castrate testosterone levels; molecular characterization of the resistant state, including assessment of AR and MYC pathway activity, remains to be established. In addition, the duration of the tumor studies was insufficient to determine whether the resistant phenotype is stable or reversible following androgen restoration. These questions can be directly addressed using the pharmacologically controlled models established in this study. A recent study reported that complete suppression of testosterone, FSH, and LH with Degarelix was more effective against AR-indifferent PC-3 and DU145 cells than against AR-dependent VCaP cells [62], raising the possibility that profound HPG-axis suppression may retain activity in tumors that are less dependent on AR signaling. Whether Degarelix has therapeutic activity as a single agent or in combination with AR-directed therapies such as enzalutamide remains poorly defined. The immunocompetent models established here provide a platform to investigate these questions while enabling evaluation of tumor-immune interactions during androgen withdrawal, resistance, and androgen restoration.

In summary, we established Degarelix treatment and withdrawal as a reversible, non-surgical model of the castration-regeneration cycle in immunocompetent mouse models. Monthly single-agent dosing achieved sustained castrate testosterone levels and reproduced the canonical prostatic response to androgen withdrawal, all of which were resolved upon drug clearance. The principal innovation is restoring endogenous androgen signaling by recovering the intact HPG axis, rather than replacing testosterone exogenously, providing a physiologically relevant model for studying androgen cycling and bipolar androgen therapy. Our innovative immunocompetent experimental models of PC reproduced the clinical trajectory of an initial response followed by acquired resistance to castration. These findings demonstrate that pharmacologic androgen deprivation is sufficient to drive resistance in an immunocompetent host. We show that the metabolic response to androgen deprivation differs between mice and men, establishing a versatile and relevant preclinical framework. The models generated in this work enable investigation of the mechanisms underlying ADT response and acquired resistance, while preserving an intact immune system. It also provides novel preclinical models to study tumor-immune interactions and the consequences of androgen re-exposure within the same experimental platform.

## Consent for publication

Not applicable.

## Availability of Data and Materials

Cell lines and models described in this study are available upon request through the Mathew Laboratory, subject to applicable UNMC material transfer agreements and institutional approvals. The software and algorithms for data analyses used in this study are published and referenced throughout the methods section. This study does not report original code. Any additional information required to reanalyze the data reported in this paper is available from the lead contact upon request.

## Funding

This work was supported by the National Institutes of Health/National Cancer Institute, NIGMS COBRE (P20GM121316), R01CA274605, American Cancer Society Institutional Research Grant, and FPBCC pilot grants to G. Mathew, and the APC was funded by the same awards. L.M. Cook was supported by an American Cancer Society Research Scholar Grant (RSG-19-127-01-CSM) and NCI R01CA274605.

We acknowledge the use of 1. The In Vivo Imaging Spectrum core facility is supported by Cancer Center Support Grant P30 CA036727 and COBRE P20 GM121316. 2. Tissue Sciences Facility, RRID: SCR_012465, which receives support from the Cancer Center Support Grant P30 CA036727 and the UNMC Department of Pathology, Microbiology, and Immunology. Research reported in this publication was also supported by the National Cancer Institute of the National Institutes of Health under award number P30 CA036727 (Cancer Center Support Grant). The content is solely the responsibility of the authors and does not necessarily represent the official views of the National Institutes of Health.

The authors used Grammarly to improve the quality of the writing for specific sections of the manuscript and BioRender.com to produce the experimental timeline schematics included in specific figures.

## Authorship

Conceptualization, A.R., D.C.G, L.M.C, and G.M.; methodology, A.R., D.C.G, G.W., and G.M.; investigations, A.R., D.C.G, and G.M.; formal analyses, A.R., D.C.G, G.W., and G.M. data curation, A.R., D.C.G, G.W., and G.M.; resources, A.R., D.C.G, G.W., L.M.C; writing, A.R., D.C.G, and G.M.; funding acquisition, L.M.C, and G.M. The contributions (manuscript writing and revisions) of the NIH author (L.M.C.) were made as part of their official duties as NIH federal employees, are in compliance with agency policy requirements, and are considered Works of the United States Government. However, the findings and conclusions presented in this paper are those of the authors and do not necessarily reflect the views of the NIH or the U.S. Department of Health and Human Services.

## Acknowledgments

The authors thank all members of the Cook and Mathew laboratories for helpful discussions and critical reading of the manuscript.

## Supplementary information

**Supplemental Figure 1**: H&E staining of the urogenital system shows prostate involution following Degarelix treatment and prostate regeneration following Degarelix withdrawal. **(A)** Bladder and prostate lobes; normal prostate glandular architecture was observed in the vehicle-treated cohort. **(B)** Chemical castration with Degarelix results in reduced prostate size. **(C)** Staining of the urogenital system following Degarelix withdrawal revealed prostate regeneration, with glandular architecture comparable to that of untreated controls. Images: Magnification = 20x, scale bar = 2mm. Images shown from the 8-week time point. ‘B’ indicates bladder; ‘AP’ indicates anterior prostate.

**Supplemental Figure 2**: Masson’s Trichrome staining of the prostate revealed stromal thickening following castration, which reverted to a single stromal layer following prostate regeneration. **(A)** Masson’s Trichrome staining of prostate tissue from the vehicle-treated cohort showed normal architecture with a single stromal tissue layer. **(B)** Degarelix-induced chemical castration resulted in stromal thickening. **(C)** After Degarelix withdrawal, Masson’s Trichrome staining revealed prostate gland regeneration and stromal thinning comparable to vehicle-treated controls. **(D)** Quantification of stromal thickness showed a significant increase after castration, with no significant difference after prostate regeneration compared with vehicle-treated controls. Images shown from the 8-week time point. Images: Magnification = 20x, scale bar = 300µm. ‘AP’ indicates the anterior prostate. ImageJ was used to measure stromal thickness (n = 10 areas per group). **** represents p < 0.0001; one-way ANOVA with Dunnett’s multiple comparisons.

