## Supplementary figures for "An Immunocompetent Mouse Model of Degarelix Induced Chemical Castration to Investigate Acquired Resistance to Androgen Deprivation"

#### Supplemental Figure 1

H&E of the Urogenital System

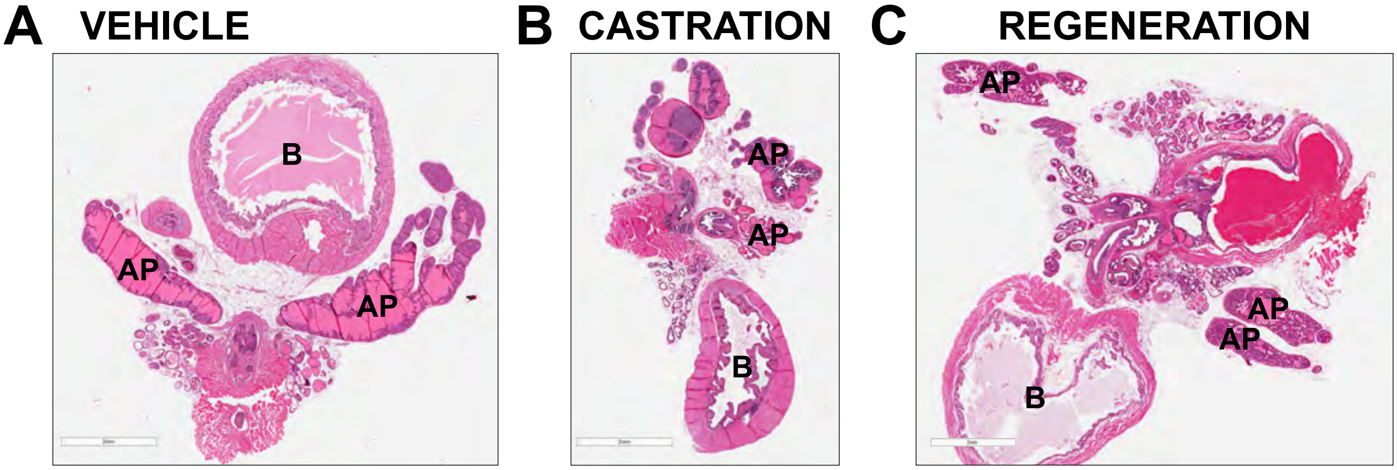

### Supplemental Figure 2

Masson's Trichrome Staining of the Urogenital System

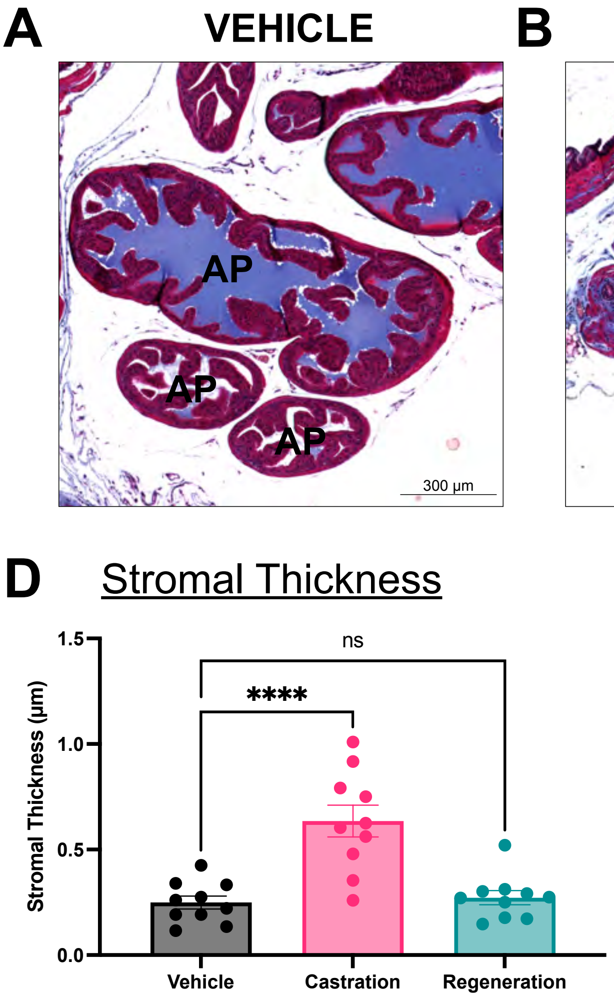
